# Adolescent alcohol exposure disrupts retrosplenial cortex physiology in adult males

**DOI:** 10.1101/2025.09.29.679287

**Authors:** Lisa R Taxier, Lili S Kooyman, Sloan Y Markowitz, Jessica A Wojick, Thomas L Kash

## Abstract

Adolescent binge drinking can lead to a myriad of issues in adulthood, including psychiatric comorbidities that worsen clinical prognosis. Efforts to understand the effects of adolescent alcohol exposure on later health outcomes have unveiled lasting effects on physiology of brain regions critical for associative learning processes, including the hippocampus and prefrontal cortex (PFC). Here, we tested the hypothesis that adolescent alcohol exposure would produce lasting alterations in physiology within the retrosplenial cortex (RSC), another region critical for associative learning that is reciprocally connected to the hippocampus and PFC. In support of this hypothesis, we found that adolescent intermittent ethanol (AIE) vapor exposure resulted in lasting reductions in intrinsic excitability in RSC pyramidal cells of adult mice. Importantly, these changes were sex-specific, occurring in males but not females. This work suggests that the RSC may be a key, vulnerable locus to adolescent alcohol’s detrimental effects.

**NEW & NOTEWORTHY:** This study is the first to demonstrate that adolescent alcohol exposure produces lasting alterations in adult retrosplenial cortex physiology, providing novel insight into how adolescent alcohol exerts detrimental effects on adult brain function.

## INTRODUCTION

Effects of adolescent alcohol exposure on adult cognition have been well documented, with a particular focus on vulnerability of the prefrontal cortex (PFC)(1–4) and the hippocampus(5,6). Behavioral consequences stemming from adolescent alcohol use include disordered memory, executive function, and behavioral flexibility (7–9). These changes often persist through adulthood, suggesting that the adolescent brain is uniquely susceptible to alcohol exposure. Alterations in hippocampus and prefrontal cortex-dependent behaviors following adolescent alcohol exposure are accompanied by disruptions to excitability and physiology in these regions. For example, adolescent alcohol exposure resulted in higher likelihood of long term potentiation following low theta burst stimulation in the adult rat hippocampus compared to alcohol-free controls (10), and increased amplitude of NMDA receptor-mediated ePSCs in hippocampal pyramidal cells (11). In the prelimbic cortex, adolescent alcohol exposure altered intrinsic excitability of pyramidal neurons (12) and increased excitability of somatostatin interneurons (4). Importantly, as with behavioral dysfunction, demonstrated effects of adolescent alcohol exposure on physiology often persist into abstinent adulthood in rodents, suggesting that exposure to alcohol during the adolescent window fundamentally alters typical developmental trajectories.

Although much literature points towards the dangerous impact of adolescent alcohol exposure on the PFC and hippocampus, other structures have received comparatively little attention. The retrosplenial cortex (RSC), which is reciprocally connected to the hippocampus and PFC(13) and plays a key role in associative learning, is another structure that may be vulnerable to alcohol. For instance, functional connectivity of the RSC is modulated by alcohol exposure in humans(14) and mice(15). In adults with AUD, the RSC is active in response to alcohol-associated cues(16,17). This finding is mirrored in adolescents with AUD, suggesting that the RSC responds to alcohol during brain maturation(18). Adult rats that underwent adolescent intermittent ethanol treatment exhibit increased cortical thickness of RSC(19); yet, the functional relevance of this lasting change in morphology is unknown. Importantly, cells within the RSC exhibit unique electrophysiological phenotypes that emerge across the adolescent window (20), suggesting that this region undergoes developmentally-regulated changes during this time. Although persistent physiological dysfunction in other regions critical for associative learning has been linked to adolescent alcohol exposure, whether the RSC is similarly vulnerable to adolescent alcohol is unknown. Indeed, despite its extensive connectivity to brain regions that are susceptible to alcohol during development and adulthood, no work has been done to explore functional changes within the RSC that result from adolescent alcohol exposure.

Here, we explored whether binge-like exposure to alcohol during adolescence via the adolescent intermittent ethanol (AIE) vapor model leads to disrupted RSC physiology in adulthood. Our results show that in male, but not female mice, AIE produces persistent effects on physiology of adult RSC pyramidal cells. In particular, we observed diminished intrinsic excitability of RSC pyramidal neurons in adult male, but not female mice exposed to AIE, as well as a reduction in excitatory drive in the adult male RSC. We also found that AIE reduced the number of local excitatory synaptic inputs to adult male RSC pyramidal neurons. Combined, these data implicate the RSC as a novel site for sex-specific and persistent effects of adolescent alcohol.

## MATERIALS AND METHODS

### Animals

Male and female C57Bl/6J mice (n = 32); Stock #: 000664, Jackson Laboratories) were obtained from Jackson Laboratories at postnatal day 21 (P21). Mice were group-housed 4/cage and maintained on a 12-hour reverse light-dark cycle in temperature- and humidity-controlled facilities, with *ad libitum* access to food (Prolab Isopro RMH 3000, LabDiet) and water. All procedures followed the National Institutes of Health Guide for the Care and Use of Laboratory Animals and were approved by the University of North Carolina-Chapel Hill School of Medicine Institutional Animal Care and Use Committee.

### Adolescent Intermittent Ethanol

Mice were exposed to air or AIE via air or ethanol vapor chamber on a 2 days-on, 2 days-off schedule, for 16 hrs/day beginning at P28, and ending at P60. Vapor chambers were calibrated to produce average blood ethanol concentrations (BECs) meeting the National Institute on Alcohol Abuse and Alcoholism definition of binge alcohol exposure (0.08%, or 0.08 grams of alcohol per deciliter or higher). Tail blood was taken at 3 interspersed intervals throughout the duration of AIE, and BECs were measured using an Analox-AM1 (Analox Technologies).

### Slice electrophysiology

Whole-cell patch-clamp recordings were obtained from pyramidal cells residing in layer 5/6 of the anterior granular RSC (within -1.06 to -1.7 AP in stereotaxic coordinates). Following rapid decapitation, brains were extracted and immersed in a chilled sucrose cutting solution (194 mM sucrose, 20 mM NaCl, 4.4 mM KCl, 2 mM CaCl_2_, 1 mM MgCl_2_, 1.2 mM NaH_2_PO_4_, 10 mM D-glucose, and 26 mM NaHCO) oxygenated with 95% O_2_/5% CO_2_. Coronal sections (250 μM) containing the RSC were obtained using a Compresstome VF-510-OZ (Precisionary Instruments) and allowed to recover for 30 minutes in 35°C oxygenated artificial cerebrospinal fluid (aCSF; 124 mM NaCl, 4.4 mM KCl, 1 mM NaH2PO4, 1.2 mM MgSO4, 10 mM D-glucose, 2 mM CaCl2, and 26 mM NaHCO3) prior to the start of recording. A cesium methanesulfonate-based internal solution (135 mM cesium methanesulfonate, 10 mM KCl, 1 mM MgCl2, 0.2 mM EGTA, 4 mM MgATP, 0.3 mM Na2GTP, 20 mM phosphocreatine, with 1mg/mL QX-314) was used during recordings of spontaneous inhibitory and excitatory post-synaptic currents (sIPSCs and sEPSCs), and a potassium gluconate-based internal solution (135 mM K-gluconate, 5 mM NaCl, 2 mM MgCl2, 10 mM HEPES, 0.6 mM EGTA, 4 mM Na2ATP, 0.4 mM Na2GTP) was used for intrinsic excitability experiments. Recording pipettes (2–4 MΩ) were pulled from thin-walled borosilicate glass capillaries using a P95 pipette puller (Sutter Instruments). Signals were acquired using an Axon Multiclamp 700B amplifier (Molecular Devices), digitized at 10 kHz, filtered at 3 kHz, and analyzed in Easy Electrophysiology. Series resistance (*R*_s_) was monitored without compensation, and changes in *R*_s_ exceeding 20% were used as exclusion criteria.

### Synaptic input mapping

Slices were prepared as described above. Synaptic input mapping was conducted with Laser Assisted Stimulation and Uncaging software (LASU, Scientifica) as previously published (21). 35 mL RuBi-glutamate (300 μM) in aCSF was oxygenated and recirculated, and was photolysed using a pulsed 405 nm laser beam (50 mw) at 5x magnification across a stimulus grid (200 μM spacing) placed across the RSC, including all cortical layers. Uncaging at each stimulation site was guided by mirror galvanometers controlled by LASU software, and was separated by 5s intervals. To isolate IPSCs and EPSCs, maps were recorded at a holding potential of +10 mV or -55 mV, respectively, and a cesium methanesulfonate internal solution (135 mM cesium methanesulfonate, 10 mM KCl, 1 mM MgCl2, 0.2 mM EGTA, 4 mM MgATP, 0.3 mM Na2GTP, 20 mM phosphocreatine, with 1mg/mL QX-314) was used. IPSCs were observed within a 2–50 ms window after the stimulus for each sweep, whereas EPSCs were observed within a 3-25 ms window (Brill et al, 2016). Amplitude of each uncaging-evoked event was recorded in pClamp 10.7, and individual synaptic input maps were combined to generate averaged synaptic input maps by recording the amplitude, in pA, and x and y distance, in microns, of each responsive stimulation site from the soma. These values were then binned at 200 μm intervals. Smooth contours were derived in MATLAB using linear interpolation between 200 μm bins.

### Statistical analyses

Statistical analyses were performed using GraphPad Prism 10 software. Unpaired *t* tests were used to compare total number of synaptic input sites, rheobase, action potential (AP) kinetic metrics, and differences in amplitude/frequency of spontaneous excitatory or inhibitory currents. Two-way ANOVAs using treatment (Air vs AIE) and distance to soma as independent variables were used for synaptic input mapping experiments to compare summed cumulative synaptic inputs. Significant ANOVA interactions were followed by Šídák’s multiple-comparisons test, with significance set at *p*□<□0.05.

## RESULTS

### AIE produces diminished intrinsic excitability in adult RSC of males, but not females

We first examined whether the excitability of cells within the adult RSC was altered by AIE. We exposed mice to air or AIE from P28-P60, and allowed mice to age alcohol-free until P90, when we collected slices containing the anterior RSC for whole-cell patch-clamp electrophysiology. In males that underwent AIE, rheobase, or minimum amount of current required for an action potential to fire, was significantly higher compared to air controls (Fig 1A and B; *t*_(16)_ = 3.07, *p* < 0.01). This effect was sex-specific, as rheobase in females was unaltered following AIE (Fig 1E and F; *t*_(19)_ = 0.76, *p* = 0.46). We next examined action potentials fired in response to increasing stepwise current injections, and found that in AIE-exposed males, cells fired fewer action potentials in response to increased current injections compared to air controls (Fig 1C and D; two-way repeated measures ANOVA, *F*_(1,16)_ = 8.58, AIE: *p* <0.01). There was also a significant interaction between current step and treatment (*F*_(20,320)_ = 3.07, *p* < 0.0001). As with rheobase, this effect of AIE was sex-specific; there were no differences in action potentials fired in response to increasing current injections in female AIE-exposed adults compared to air controls (Fig 1E and F; *F*_(1, 17)_ = 0.02, *p* = 0.89), nor an interaction between current step and treatment (*F*_(20, 340)_ = 0.71, *p* = 0.81). Combined, these data suggest that AIE diminishes intrinsic excitability in adult male, but not adult female RSC.

**Figure 1.**
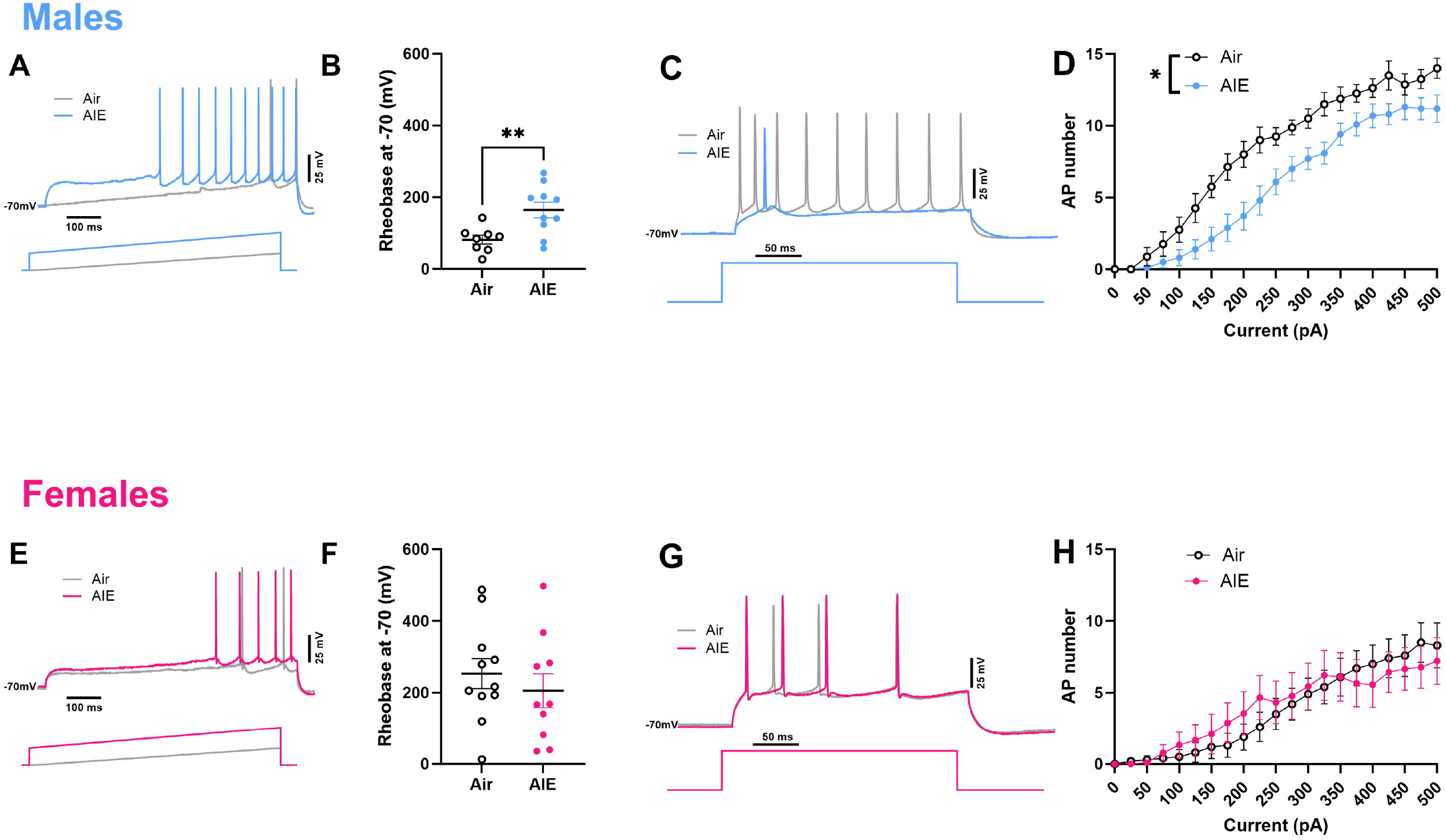
AIE diminishes RSC pyramidal cell intrinsic excitability in adult males, but not females. Representative traces from males (A) and females (E) show examples of increased rheobase in cells from AIE-exposed males (B) but no AIE-mediated effects on rheobase in females (F). Representative traces from males (C) and females (G) show examples of current-induced firing, which is reduced in cells from AIE-exposed males (D), but not in females (H). n = 8 cells, Air males; 10 cells, AIE males; 11 cells, Air females, 11 cells, AIE females; * p < 0.05, ** p <0.01, error bars indicate SEM.

Intrinsic excitability, which is a crucial regulator of learning and memory processes, can be modulated by changes in ion channel distribution and function(22). To test the hypothesis that AIE exerts effects on intrinsic excitability via altering ion channel function, we assessed metrics of action potential (AP) kinetics including AP latency, height, threshold, and half-width. AIE increased AP latency in males (Fig 2A, *t*_(16)_ = 2.76, *p* = 0.01), but had no significant effect on AP height (Fig 2B), threshold (Fig 2C), or half-width (Fig 2D). Again, any observed effects of AIE on AP kinetics were sex-specific; AIE had no demonstrable effects on AP kinetics in adult female RSC (Fig 2E-H). Given that AIE increased AP latency in males, taken together with observed effects of AIE on intrinsic excitability, these data suggest that diminished intrinsic excitability may be due in part to AIE modulation of ion channel function within the adult male RSC.

**Figure 2.**
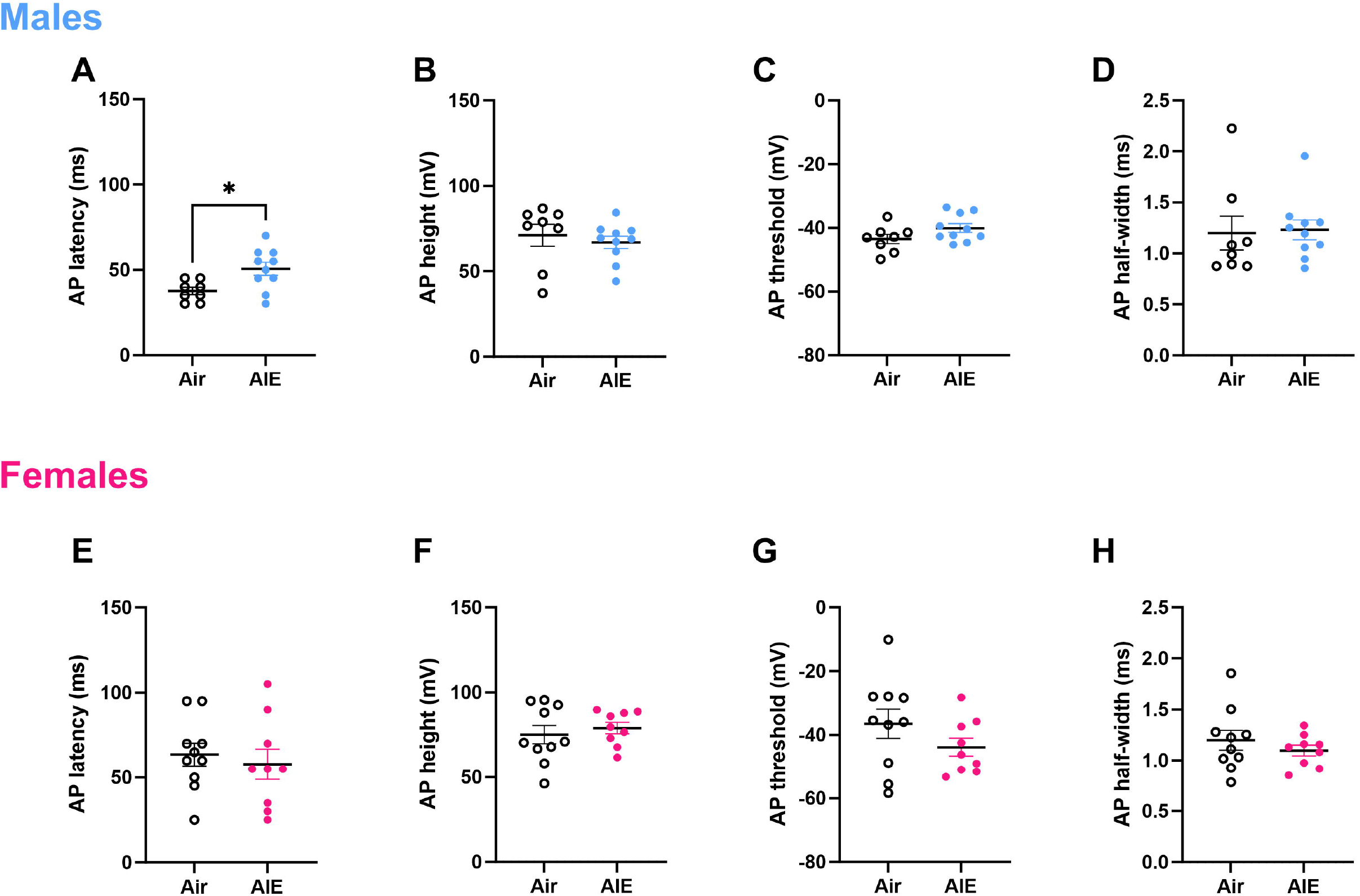
AIE increases AP latency in RSC pyramidal cells of adult males, but not females. Latency to fire an action potential is increased in RSC pyramidal cells from adult males (A) but not females (E). AP height (B, F), AP threshold (C, G) and AP half-width (D, H), were unchanged by AIE in both sexes. N = 8 cells, Air males; 10 cells, AIE males; 11 cells, Air females, 11 cells, AIE females; * p < 0.05, error bars indicate SEM.

### AIE reduces excitatory drive in adult RSC of males, but not females

To evaluate spontaneous synaptic transmission in adult RSC following AIE, we used a cesium methanesulfonate-based internal solution (see methods) to isolate sEPSCs and sIPSCs in the same cells while voltage clamping at -55 mV and +10 mV, respectively (Fig 3A, F). This approach allows for the evaluation of excitatory-inhibitory (E/I) drive onto individual neurons. Males displayed a trend towards AIE-diminished frequency of sEPSCs (unpaired *t*-test, *t*_(24)_ = 1.74, *p* = 0.09), and an impact of AIE on sEPSC amplitude such that RSC cells from AIE-exposed males had a significantly lower amplitude than those from air-exposed controls (*t*_(24)_ = 2.11, *p* = 0.05). There was no such effect of AIE on sIPSC frequency (Fig 3D) or amplitude (Fig 3E) in males. In females, there were no significant differences in frequency or amplitude of sEPSCs (Fig 3G, H); this null effect was also present for both frequency and amplitude of sIPSCs (Fig 3I,J).

**Figure 3.**
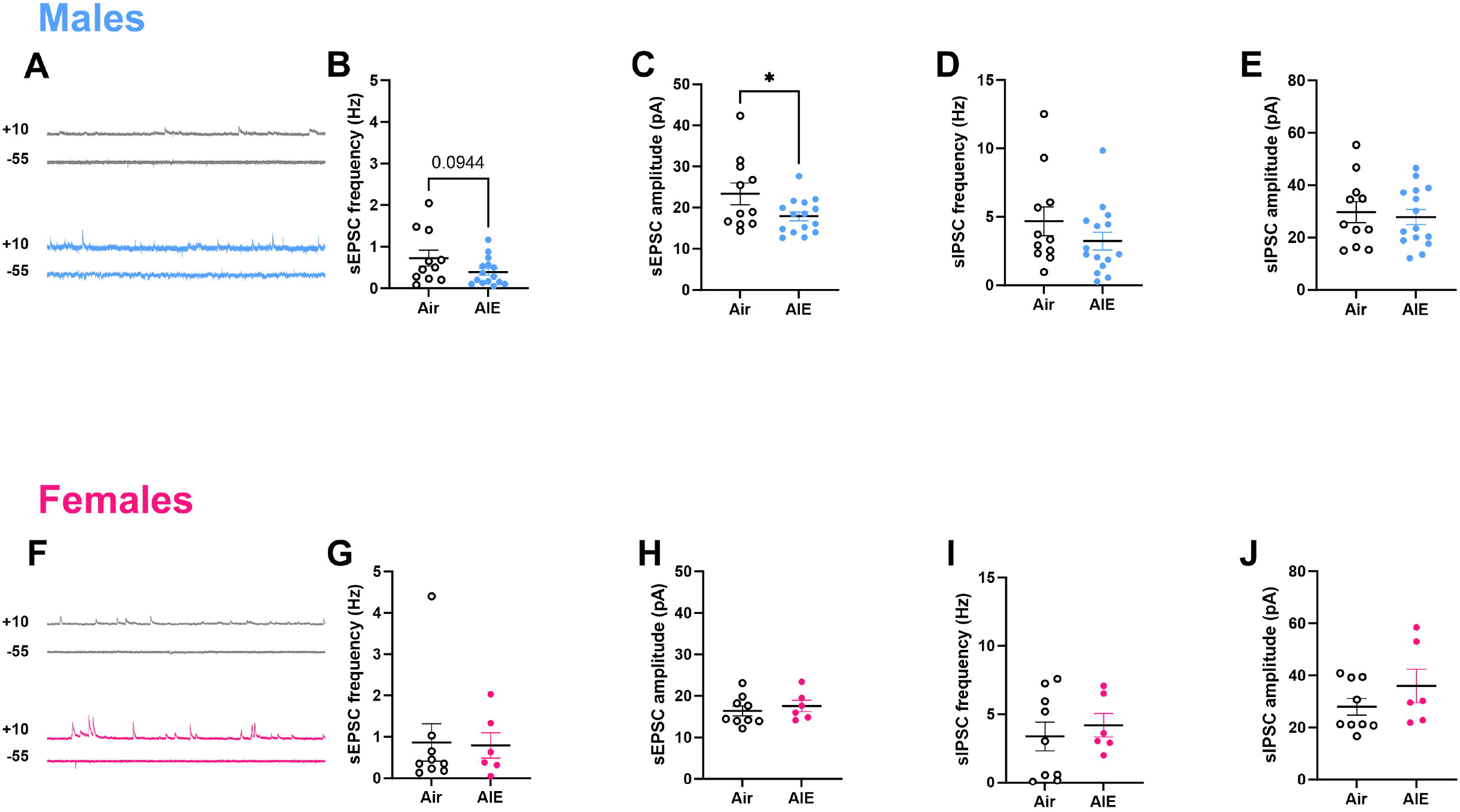
AIE reduces spontaneous excitatory drive in adult RSC of males, but not females. Representative recordings from Males (A) and Females (F) showing spontaneous firing of EPSCs recorded at -55 mV, or IPSCs recorded at +10 mV. sEPSC frequency (B) and amplitude (C) were reduced in cells from AIE-exposed males, but not females (G, H). sIPSC frequency (D,I) and amplitude (E,J) were unaffected by AIE exposure in either sex. n = 11 cells, Air males, 15 cells, AIE males; 9 cells, Air females, 6 cells, AIE females; *p <0.04, error bars indicate SEM.

### AIE reduces the number of local excitatory synaptic inputs to adult male RSC pyramidal neurons

Excitatory neurons within the superficial layers of RSC, which receive robust local inhibitory input from neighboring interneurons, project through deeper layers of RSC and into the corpus callosum(23). Diminished intrinsic excitability and reduced glutamatergic drive onto layer 5/6 pyramidal neurons of male RSC following AIE could potentially arise from decreased excitatory or increased inhibitory input from layer 2/3, or from fast spiking interneurons present across the RSC laminar extent. To examine whether local synaptic input might contribute to our observed effect of AIE on intrinsic excitability, we conducted synaptic input mapping experiments in adult RSC following AIE. We patched onto pyramidal cells in layer 5/6, and uncaged Rubi-glutamate using focal laser application at 200μM intervals across the extent of the RSC (Fig. 4A) while voltage clamping at -55 and +10 to isolate uncaging-evoked EPSCs and IPSCs, respectively. This approach allowed for the detection of the presence, strength, and spatial location of local synaptic excitatory and inhibitory inputs to patched cells, and we generated averaged synaptic input maps to all patched cells to visualize the spatial pattern and strength of such inputs (Fig. 4B and C). In males, the total number of sites where EPSCs, but not IPSCs, were evoked was reduced in cells from AIE-exposed mice relative to air-exposed controls (Fig. 4D, left; *t*_(16)_ = 2.34, *p* = 0.03). In line with this, when cumulative amplitude was collapsed across the interlaminar extent (i.e., horizontal distance to soma), two-way ANOVA revealed a trend towards an effect of AIE (Fig 4E, left; *F*_(1,160)_□=□2.85; *p*□=□0.09), such that the amplitude of uncaging-elicited EPSCs was lower in cells from AIE-exposed males compared with air-exposed controls. Similarly, when cumulative amplitude was collapsed across the intralaminar extent (i.e., vertical distance to soma), two-way ANOVA revealed a trend towards an effect of AIE (Fig 4E, right; *F*_(1,162)_ = 3.02, *p* = 0.08), such that amplitude of uncaging-evoked EPSCs was lower in cells from AIE-exposed males compared with air-exposed controls. For IPSCs, there was no significant effect when cumulative amplitude was collapsed across either the intralaminar (Fig 4F, left) or the interlaminar (Fig 4F, right) extent. In females, as with every other examined metric, there was no significant effect of AIE on total input sites for EPSCs or IPSCs (Fig 4G), no significant effect of AIE on EPSC amplitude along the dorsal/ventral axis (Fig 4H, left) or medial/lateral axis (Fig 4H, right), and no significant effect of AIE on IPSCs amplitude along the dorsal/ventral axis (Fig 4I, left) or medial/lateral axis (Fig 4I, right). For all collapsed cumulative amplitude measures, regardless of sex, there was a significant effect of either dorsal/ventral or medial lateral distance such that amplitude of evoked EPSCs and IPSCs was higher at uncaging sites near the soma.

**Figure 4.**
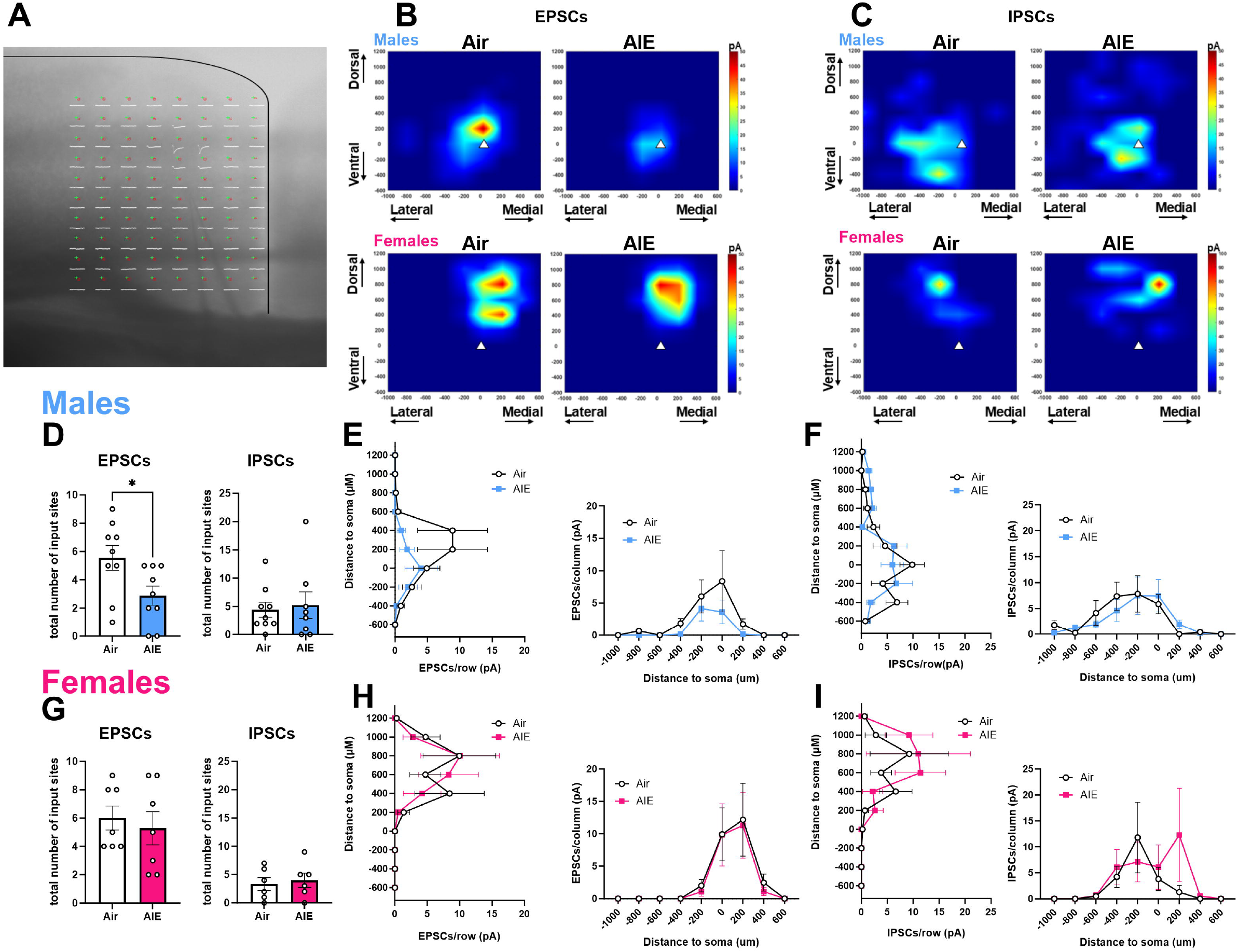
AIE reduces the number of local excitatory synaptic inputs to adult male RSC pyramidal neurons. (A) Representative grid superimposed over the extent of the left hemisphere of the anterior RSC, with corresponding traces in white detailing sEPSCs elicited by glutamate uncaging at each grid site (green crosses). (B) Representative summed input maps of sEPSCs elicited by glutamate uncaging in males (top) and females (bottom), in RSC pyramidal cells from both Air (left) and AIE (right)-exposed mice. (C) Representative summed input maps of sIPSCs elicited by glutamate uncaging in males (top) and females (bottom), in RSC pyramidal cells from both Air (left) and AIE (right)-exposed mice. (D) Total number of input sites where EPSCs, but not IPSCs were elicited was reduced cells from AIE-exposed mice compared to cells from air-exposed controls. (E) Amplitude of uncaging-evoked EPSCs tended to be lower onto cells from AIE-exposed males across both the inter- and intralaminar extent, whereas amplitude of uncaging-evoked IPSCs (F) was unaffected by AIE. (G) In females, there was no AIE-mediated difference in total number of input sites where EPSCs or IPSCs were evoked by glutamate uncaging, nor was there an effect of AIE on averaged inter- or intralaminar amplitude of evoked EPSCs (H) or IPSCs (I).

## DISCUSSION

Although considerable evidence points to a role for adolescent alcohol in modulating adult behavior and brain function, questions remain about region specific mechanisms that drive such effects. Here, we provide novel evidence that the RSC is susceptible to adolescent alcohol exposure in a sex-specific manner. We show that AIE produces lasting decreases in intrinsic excitability of pyramidal cells within the adult male, but not female RSC, and that such changes in intrinsic excitability are accompanied by increased action potential latency. We also demonstrate that AIE produces reductions in spontaneous excitatory drive onto adult male, but not female, RSC pyramidal cells, and that AIE produces diminished local excitatory synaptic input onto adult male, but not female RSC pyramidal cells.

One surprising result of the current experiments was that there was no detectable effect of AIE on RSC physiology in females. One possibility for lack of observed effect in females is the timing of exposure; although exposure across P28-P55 captures broadly agreed-upon definitions of the adolescent window(24), it may be the case that female mice may be more vulnerable to alcohol exposure at earlier or later time points than were investigated in the current study. Another possibility is that females are more resilient to the effects of alcohol exposure during the adolescent window than males, which presents an intriguing possibility that deserves additional exploration. Although we did not make a statistical comparison between sexes, we observed that females seem to exhibit a much less excitable phenotype at baseline compared to males. This observation highlights the need for future work to investigate baseline sex differences in electrophysiological phenotypes within the RSC. Interestingly, sex differences in adult behavior following exposure to alcohol during adolescence are well documented (7,9,25); differences in RSC electrophysiological phenotype following AIE may be one mechanism underlying behavioral differences, although this remains to be directly tested.

Our finding that adolescent alcohol robustly and persistently diminishes intrinsic excitability within the adult male RSC is particularly striking given studies demonstrating the opposite effect of adolescent alcohol on intrinsic excitability of pyramidal neurons within other structures such as the prelimbic cortex (2,26). Furthermore, in the hippocampus, AIE facilitates low stimulus intensity-induced LTP (27), suggesting a shift towards AIE-driven excitation within this region. Although one popular theory posits that adolescent-like phenotypes favoring excitation over inhibition persist into adulthood following adolescent alcohol exposure, our data indicate that the RSC may be an exception to this idea (28). However, it may also be the case that the RSC maintains a different profile of excitatory and inhibitory balance during adolescence than other cortical structures. Indeed, one study suggests that neurons within the RSC that exhibit an afterdepolarization following action potential firing do not emerge until later in development, which is notable given that this particular firing property seems to enhance excitability (20). Although we did not examine intrinsic excitability in the adolescent RSC in the absence of alcohol exposure, future work should assess whether adolescent alcohol perpetuates existing adolescent-like phenotypes within this region.

Surprisingly, we found no effect of adolescent alcohol on spontaneous inhibitory drive or on local inhibitory synaptic input to adult RSC pyramidal cells. Because the RSC is populated with several interneuron subtypes including parvalbumin-expressing interneurons, vasoactive inhibitory peptide-expressing cells, and somatostatin-expressing interneurons, we suspected that diminished intrinsic excitability of pyramidal cells within adult RSC following AIE may be due to increased inhibition from these local cell populations. However, given our synaptic input mapping findings and lack of effect of AIE on spontaneous inhibitory drive, we speculate that long-range inhibitory projections to RSC, such as those from the PFC or the hippocampus, may play a more consequential role in driving the effects of AIE on decreased intrinsic excitability. For example, a population of somatostatin neurons in the PFC project to the RSC(29) and excitability of prelimbic somatostatin neurons is increased following adolescent alcohol exposure(4). This suggests the possibility that following AIE, long-range somatostatin projections from the PFC impinge upon RSC pyramidal cells to inhibit glutamatergic drive and diminish intrinsic excitability.

Taken together, our data demonstrate that binge-like exposure to alcohol during adolescence has lasting effects on physiological function of the adult male RSC. These findings add critical information about how adolescent alcohol can shape adult brain function, and point towards a novel brain target for additional study. Given the dense interconnectedness of the RSC, future studies should examine the role of AIE on both inputs and outputs to RSC. Such research will ultimately add to mechanistic understanding of how AIE might produce lasting adult dysfunction and ultimately lead to the development of new therapeutics.

## DATA AVAILABILITY

Data from the present experiments are available from the corresponding author upon reasonable request.

## AUTHOR CONTRIBUTIONS

LRT and TLK designed experiments. LRT and LSK conducted behavioral experiments. LRT conducted slice electrophysiology experiments with contributions from SYM. LRT wrote the paper with contributions from LSK, SYM, and TLK.

## REFERENCES

1. De Bellis MD, Narasimhan A, Thatcher DL, Keshavan MS, Soloff P, Clark DB. Prefrontal Cortex, Thalamus, and Cerebellar Volumes in Adolescents and Young Adults with Adolescent-Onset Alcohol Use Disorders and Comorbid Mental Disorders. Alcohol: Clinical and Experimental Research. 2005;29(9):1590–600. doi:10.1097/01.alc.0000179368.87886.76

2. Salling MC, Skelly MJ, Avegno E, Regan S, Zeric T, Nichols E, et al. Alcohol consumption during adolescence in a mouse model of binge drinking alters the intrinsic excitability and function of the prefrontal cortex through a reduction in the hyperpolarization-activated cation current. J Neurosci. 2018 Jul 4;38(27):6207–22. doi:10.1523/JNEUROSCI.0550-18.2018 PubMed PMID: 29915134.

3. Sicher AR, Duerr A, Starnes WD, Crowley NA. Adolescent Alcohol and Stress Exposure Rewires Key Cortical Neurocircuitry. Front Neurosci. 2022 May 17;16:896880. doi:10.3389/fnins.2022.896880 PubMed PMID: 35655755; PubMed Central PMCID: PMC9152326.

4. Sicher AR, Starnes WD, Griffith KR, Dao NC, Smith GC, Brockway DF, et al. Adolescent binge drinking leads to long-lasting changes in cortical microcircuits in mice. Neuropharmacology. 2023 Aug 15;234:109561. doi:10.1016/j.neuropharm.2023.109561

5. Wooden JI, Thompson KR, Guerin SP, Nawarawong NN, Nixon K. Consequences of Adolescent Alcohol Use on Adult Hippocampal Neurogenesis and Hippocampal Integrity. Int Rev Neurobiol. 2021;160:281–304. doi:10.1016/bs.irn.2021.08.005 PubMed PMID: 34696876; PubMed Central PMCID: PMC8878345.

6. De Bellis MD, Clark DB, Beers SR, Soloff PH, Boring AM, Hall J, et al. Hippocampal Volume in Adolescent-Onset Alcohol Use Disorders. AJP. 2000 May;157(5):737–44. doi:10.1176/appi.ajp.157.5.737

7. Crews FT, Vetreno RP, Broadwater MA, Robinson DL. Adolescent Alcohol Exposure Persistently Impacts Adult Neurobiology and Behavior. Pharmacol Rev. 2016 Oct;68(4):1074–109. doi:10.1124/pr.115.012138 PubMed PMID: 27677720; PubMed Central PMCID: PMC5050442.

8. Bergstrom HC, McDonald CG, Smith RF. Alcohol exposure during adolescence impairs auditory fear conditioning in adult Long-Evans rats. Physiology & Behavior. 2006 Jul 30;88(4):466–72. doi:10.1016/j.physbeh.2006.04.021

9. Chandler LJ, Vaughan DT, Gass JT. Adolescent alcohol exposure results in sex-specific alterations in conditioned fear learning and memory in adulthood. Front Pharmacol. 2022 Feb 8;13:837657. doi:10.3389/fphar.2022.837657 PubMed PMID: 35211024; PubMed Central PMCID: PMC8861326.

10. Risher ML, Fleming RL, Risher C, Miller KM, Klein RC, Wills T, et al. Adolescent Intermittent Alcohol Exposure: Persistence of Structural and Functional Hippocampal Abnormalities into Adulthood. Alcohol Clin Exp Res. 2015 Jun;39(6):989–97. doi:10.1111/acer.12725 PubMed PMID: 25916839; PubMed Central PMCID: PMC4452443.

11. Swartzwelder HS, Park MH, Acheson S. Adolescent Ethanol Exposure Enhances NMDA Receptor-Mediated Currents in Hippocampal Neurons: Reversal by Gabapentin. Sci Rep. 2017 Oct 13;7(1):13133. doi:10.1038/s41598-017-12956-6

12. Salling MC, Skelly MJ, Avegno E, Regan S, Zeric T, Nichols E, et al. Alcohol Consumption during Adolescence in a Mouse Model of Binge Drinking Alters the Intrinsic Excitability and Function of the Prefrontal Cortex through a Reduction in the Hyperpolarization-Activated Cation Current. J Neurosci. 2018 Jul 4;38(27):6207–22. doi:10.1523/JNEUROSCI.0550-18.2018 PubMed PMID: 29915134; PubMed Central PMCID: PMC6031577.

13. Vann SD, Aggleton JP, Maguire EA. What does the retrosplenial cortex do? Nat Rev Neurosci. 2009 Nov;10(11):11. doi:10.1038/nrn2733

14. Song Z, Chen J, Wen Z, Zhang L. Abnormal functional connectivity and effective connectivity between the default mode network and attention networks in patients with alcohol-use disorder. Acta Radiol. 2021 Feb 1;62(2):251–9. doi:10.1177/0284185120923270

15. Degiorgis L, Arefin TM, Ben-Hamida S, Noblet V, Antal C, Bienert T, et al. Translational Structural and Functional Signatures of Chronic Alcohol Effects in Mice. Biological Psychiatry. 2022 Jun 15;Stress, Distress, and Substance Use91(12):1039–50. doi:10.1016/j.biopsych.2022.02.013

16. Zhornitsky S, Zhang S, Ide JS, Chao HH, Wang W, Le T, et al. Alcohol expectancy and cerebral responses to cue-elicited craving in adult non-dependent drinkers. Biol Psychiatry Cogn Neurosci Neuroimaging. 2019 May;4(5):493–504. doi:10.1016/j.bpsc.2018.11.012 PubMed PMID: 30711509; PubMed Central PMCID: PMC6500759.

17. Bragulat V, Dzemidzic M, Talavage T, Davidson D, O’Connor SJ, Kareken DA. Alcohol Sensitizes Cerebral Responses to the Odors of Alcoholic Drinks: An fMRI Study. Alcohol Clin Exp Res. 2008 Jul;32(7):1124–34. doi:10.1111/j.1530-0277.2008.00693.x PubMed PMID: 18540915; PubMed Central PMCID: PMC2905852.

18. Tapert SF, Cheung EH, Brown GG, Frank LR, Paulus MP, Schweinsburg AD, et al. Neural Response to Alcohol Stimuli in Adolescents With Alcohol Use Disorder. Archives of General Psychiatry. 2003 Jul 1;60(7):727–35. doi:10.1001/archpsyc.60.7.727

19. Vetreno RP, Yaxley R, Paniagua B, Johnson GA, Crews FT. Adult rat cortical thickness changes across age and following adolescent intermittent ethanol treatment. Addiction Biology. 2017;22(3):712–23. doi:10.1111/adb.12364

20. Yousuf H, Nye AN, Moyer JR. Heterogeneity of neuronal firing type and morphology in retrosplenial cortex of male F344 rats. J Neurophysiol. 2020 May 1;123(5):1849–63. doi:10.1152/jn.00577.2019 PubMed PMID: 32267193.

21. Taxier LR, Neira S, Flanigan ME, Haun HL, Eberle MR, Kooyman LS, et al. Retrieval of an ethanol-conditioned taste aversion promotes GABAergic plasticity in the anterior insular cortex. J Neurosci. 2025 Jan 8;e0525242024. doi:10.1523/JNEUROSCI.0525-24.2024 PubMed PMID: 39779373.

22. Chen L, Cummings KA, Mau W, Zaki Y, Dong Z, Rabinowitz S, et al. The role of intrinsic excitability in the evolution of memory: Significance in memory allocation, consolidation, and updating. Neurobiology of Learning and Memory. 2020 Sep 1;173:107266. doi:10.1016/j.nlm.2020.107266

23. Brennan EKW, Sudhakar SK, Jedrasiak-Cape I, John TT, Ahmed OJ. Hyperexcitable Neurons Enable Precise and Persistent Information Encoding in the Superficial Retrosplenial Cortex. Cell Rep. 2020 Feb 4;30(5):1598-1612.e8. doi:10.1016/j.celrep.2019.12.093 PubMed PMID: 32023472.

24. Spear LP. Neurobehavioral Changes in Adolescence. Curr Dir Psychol Sci. 2000 Aug 1;9(4):111–4. doi:10.1111/1467-8721.00072

25. Kasten CR, Carzoli KL, Sharfman NM, Henderson T, Holmgren EB, Lerner MR, et al. Adolescent alcohol exposure produces sex differences in negative affect-like behavior and group I mGluR BNST plasticity. Neuropsychopharmacol. 2020 Jul;45(8):1306–15. doi:10.1038/s41386-020-0670-7

26. Galaj E, Guo C, Huang D, Ranaldi R, Ma YY. Contrasting effects of adolescent and early-adult ethanol exposure on prelimbic cortical pyramidal neurons. Drug and Alcohol Dependence. 2020 Nov 1;216:108309. doi:10.1016/j.drugalcdep.2020.108309

27. Risher ML, Fleming RL, Risher C, Miller KM, Klein RC, Wills T, et al. Adolescent Intermittent Alcohol Exposure: Persistence of Structural and Functional Hippocampal Abnormalities into Adulthood. Alcohol Clin Exp Res. 2015 Jun;39(6):989–97. doi:10.1111/acer.12725 PubMed PMID: 25916839; PubMed Central PMCID: PMC4452443.

28. Spear LP, Swartzwelder HS. Adolescent alcohol exposure and persistence of adolescent-typical phenotypes into adulthood: A mini-review. Neuroscience & Biobehavioral Reviews. 2014 Sep 1;45:1–8. doi:10.1016/j.neubiorev.2014.04.012

29. Dautan D, Monai A, Maltese F, Chang X, Molent C, Mauro D, et al. Cortico-cortical transfer of socially derived information gates emotion recognition. Nat Neurosci. 2024 Jul;27(7):1318–32. doi:10.1038/s41593-024-01647-x

